# A Diversity Outbred F1 mouse model identifies host-intrinsic genetic regulators of response to immune checkpoint inhibitors

**DOI:** 10.1101/2021.10.07.463570

**Authors:** Justin B. Hackett, James E. Glassbrook, Maria C. Muñiz, Madeline Bross, Abigail Fielder, Gregory Dyson, Nasrin Movahhedin, Jennifer McCasland, Claire McCarthy-Leo, Heather M. Gibson

## Abstract

Immune checkpoint inhibitors (ICI) have improved outcomes for a variety of malignancies; however, many patients fail to benefit. While tumor-intrinsic mechanisms are likely involved in therapy resistance, it is unclear to what extent host genetic background influences response. To investigate this, we utilized the Diversity Outbred (DO) and Collaborative Cross (CC) mouse models. DO mice are an outbred stock generated by crossbreeding 8 inbred founder strains, and CC mice are recombinant inbred mice generated from the same 8 founders. We generated 207 DOB6F1 mice representing 48 DO dams and demonstrated that these mice reliably accept the C57BL/6-syngeneic B16F0 tumor and that host genetic background influences response to ICI. Genetic linkage analysis from 142 mice identified multiple regions including one within chromosome 13 that associated with therapeutic response. We utilized 6 CC strains bearing the positive (NZO) or negative (C57BL/6) driver genotype in this locus. We found that 2/3 of predicted responder CCB6F1 crosses show reproducible ICI response. The chromosome 13 locus contains the murine prolactin family, which is a known immunomodulating cytokine associated with various autoimmune disorders. To directly test whether prolactin influences ICI response rates, we implanted inbred C57BL/6 mice with subcutaneous slow-release prolactin pellets to induce mild hyperprolactinemia. Prolactin augmented ICI response against B16F0, with increased CD8 infiltration and 5/8 mice exhibiting slowed tumor growth relative to controls. This study highlights the role of host genetics in ICI response and supports the use of F1 crosses in the DO and CC mouse populations as powerful cancer immunotherapy models.

**One Sentence Summary:** Using germline genetic analysis in Diversity Outbred and Collaborative Cross models, we identify prolactin as a host-intrinsic regulator of response to combined immune checkpoint inhibitors.

## INTRODUCTION

Cancer immunotherapy has demonstrated success against several malignancies for a subset of patients, with immune checkpoint regulators paving the way as a lead modality. Immune checkpoint molecules are a natural braking mechanism critical for balancing the activation of an immune response, while preventing excessive tissue damage or autoimmunity (*1–3*). In cancer, immune checkpoints are often co-opted by malignant cells to escape immune-mediated attack. Clinical immune checkpoint inhibitors (ICI) have thus far primarily targeted two signaling axes: Cytotoxic T-Lymphocyte-Associated protein 4 (CTLA-4) and Programmed Death 1 (PD-1) with its ligand PD-L1. Both of these signaling pathways are classified as inhibitory receptors that down-regulate CD8^+^ T cell effector functions (Reviewed in (*1–3*)). Additional checkpoint molecules are also in various stages of preclinical and clinical investigation (*4*).

Combinations of ICI therapies have also been successful in treating advanced melanoma. This approach has been extended to the treatment of lung and renal carcinomas, with evaluation against other malignancies in the pipeline (*4–10*). Despite the frequently high tumor mutational burden and associated neoantigens, the objective response rate of combined ICI in metastatic melanoma is only ~58% (*10*). Whether host-intrinsic genetic factors influence this effect remains unclear. Currently, animal models that adequately capture host-intrinsic mechanisms of resistance to ICI are lacking, potentially due to the absence of heterogeneity in inbred animal models. The human population is genetically heterogeneous, which likely contributes to the disparate response to ICI in the clinic. To better capture the dynamics of human genetic variation in an animal model where extraneous variables can be controlled we utilized the Diversity Outbred (DO) and Collaborative Cross (CC) mouse models (*11–14*). The DO mouse model (discussed below) is an extension of the CC mouse project. The CC mice are a collection of recombinant inbred (RI) mouse strains produced through ~30 generations of random outbreeding of 8 inbred founder strains (A/J, C57BL/6J, 129S1/SvImJ, NOD/LtJ, NZO/HlLtJ, CAST/EiJ, PWK/PhJ, and WSB/EiJ). This was followed by re-inbreeding, generating dozens of distinct RI CC lines. These CC strains each have fully sequenced backgrounds which can be used for genetic mapping of complex traits either for 1) discovery phase studies where multiple CC strains are tested for the phenotype of interest, or 2) validation of a locus of interest where potential driver haplotypes are known (*15*, *16*). Currently >60 CC strains are maintained by UNC Systems Genetics that are available for purchase and their online tools allow for simple exploration of founder contributions by locus.

DO mice were generated by continued outbreeding beyond the CC project, and DO mice are now maintained by The Jackson Laboratory, currently at generation 45 (~74 total including the initial CC crosses). Each DO mouse is genetically distinct, providing a model optimized for high resolution Quantitative Trait Locus (QTL) mapping that enables identification of genomic regions that influence complex traits (*15*, *16*). The Mouse Universal Genotyping Array (MUGA), now in its third iteration (GigaMUGA) genotypes 143,259 markers spanning the entire genome of each mouse (*11*). These markers are not directly used to identify causal single nucleotide polymorphisms (SNPs), but instead enable haplotype block reconstruction. Identifying the founder strain contributions across all chromosomes (Chr) allows for imputation and evaluation of all genomic SNPs. QTL effect analysis can then be performed to determine the direction and effect size of a phenotypic association to a specific founder genotype at each locus across the genome. The entirety of the data analysis pipeline is performed in the R/qtl2 package, which is optimized for use with the DO mouse population (*11*). Locus validation can then be conducted using inbred CC lines selected for haplotypes matching the QTL effects data (*12*). We have crossed inbred C57BL/6 (B6) sires with several cohorts of DO dams to produce DOB6F1 mice, which are used to investigate germline factors that contribute to ICI response.

Here we show that a Genome-Wide Association Study (GWAS) of DOB6F1 mice identifies multiple host genomic loci associated with response to ICI therapy in the B6–syngeneic B16F0 melanoma tumor model. We further validated a candidate locus in Chr13 using a panel of anticipated responder and non-responder CC mouse strains, and we identified prolactin as the candidate causal gene from within this locus. Co-administration of prolactin with ICI resulted in improved treatment response rates compared to ICI alone in inbred B6 mice.

In humans there is a single prolactin gene which is under dual transcriptional control by either the pituitary or extrapituitary promoter, with the latter allowing for expression and function external to the hypothalamic pituitary axis (*12*). In mice there are 23 members of the prolactin family clustered on Chr13 with evidence that at least some members function through the Prl receptor (*12*, *13*). Expression of the murine prolactin family genes may be regulated by tissue-dependent methylation (*12–14*). There is a significant body of research implicating prolactin in immunoregulation of both mice and humans, which describes prolactin’s modulation of B cells, macrophages, and T cells (reviewed in (*17*)). It also may have an immunomodulatory role in the setting of chronic *Trypanosoma cruzi* infection in rats (*18*, *19*). Most relevant to our study is the implication of prolactin in autoimmunity. Systemic Lupus Erythematosus (SLE) is an autoimmune condition with largely unknown etiology that results in significant damage to several organ systems (*20*). Though largely considered a “B-cell disorder” there is a robust body of work that demonstrates a significant role for T cell dysregulation in the pathogenesis of SLE (reviewed in (*21*)). Additionally, prolactin drives CD40L expression on T cells which is implicated in hyperactivation of B cells in SLE (*22*). A SNP within the human prolactin extrapituitary promoter drives elevated expression of prolactin within *in vitro*-stimulated lymphocytes, and was found to associate with SLE onset and severity (*23*). A correlation of serum prolactin levels with the onset and severity of SLE in humans is also reported (*23*, *24*). Preclinically, hyperprolactinemia drives an SLE-like condition in BALB/c mice carrying a transgene for an anti-DNA Ab heavy chain (*25*). Genetic regulation of the prolactin-driven lupus phenotype is evident, as B6 mice do not break tolerance unless they are transgenic for the anti-DNA Ab heavy chain and the lupus-like susceptibility interval *Sle3/5* (*25*). These findings are consistent with evidence of heightened self-tolerance and reduced response to immunotherapy in B6 versus BALB/c tumor models (*26*, *27*).

Collectively, we have demonstrated that the host genome influences response to ICI. Furthermore, by exploiting the power of the DO and CC models we can pinpoint host-derived regulators such as prolactin and identify actionable pathways to augment treatment response.

## RESULTS

### Response to ICI is regulated by the host genome

We and others found that inbred B6 mice bearing syngeneic B16F0 melanoma fail to respond to combined anti-CTLA-4/PD-1 ICI ((*28*) and **Fig. 1A**). To test whether genetic background influences response to ICI, we performed a pilot study with 20 DOB6F1 mice (Cohort 1, **Table 1**) which were inoculated with subcutaneous B16F0 and treated with combined anti-CTLA-4/PD-1 ICI. Once a palpable tumor was detected in all mice, treatment was initiated (days 10, 13, and 16 post-inoculation). Tumors were collected on day 19 (**Fig. 1B**). Despite inoculation with a genetically identical tumor and administration of identical treatment, these genetically distinct mice exhibited a broad range of tumor growth rate curves, suggesting that host genetic background impacts response to ICI. To compare intratumoral immune activity with tumor growth rate, we evaluated flash-frozen tumor tissue for known cytotoxic T cell signatures CD8 and IFNγ by qRT-PCR and found an inverse correlation between the expression level of both transcripts relative to final tumor volume (**Fig. 1C,D**). In a second cohort of 45 DOB6F1 mice (Cohort 2, **Table 1**), with identical treatment to Cohort 1, tumors were collected on days 17-19 as non-responding mice reached tumor burden threshold (**Fig. 1E**). Tumors from these mice showed a correlation of CD8 infiltration to final tumor burden as measured by immunohistochemistry (IHC) (**Fig. 1F**). Interestingly, we did not see a robust correlation to either density of FoxP3+ regulatory T cells or the CD8:FoxP3 ratio in the same tumor tissues (**Fig. S1A,B**). To compare tumor burden and growth rate we calculated tumor latency which we defined as the time in days until the tumor reached 150 mm^3^. Compared to treated or untreated B6 mice bearing B16F0 tumors, DOB6F1 mice had a much larger range of tumor latency and larger coefficient of variation (Fig. 1G).

**Figure 1.**
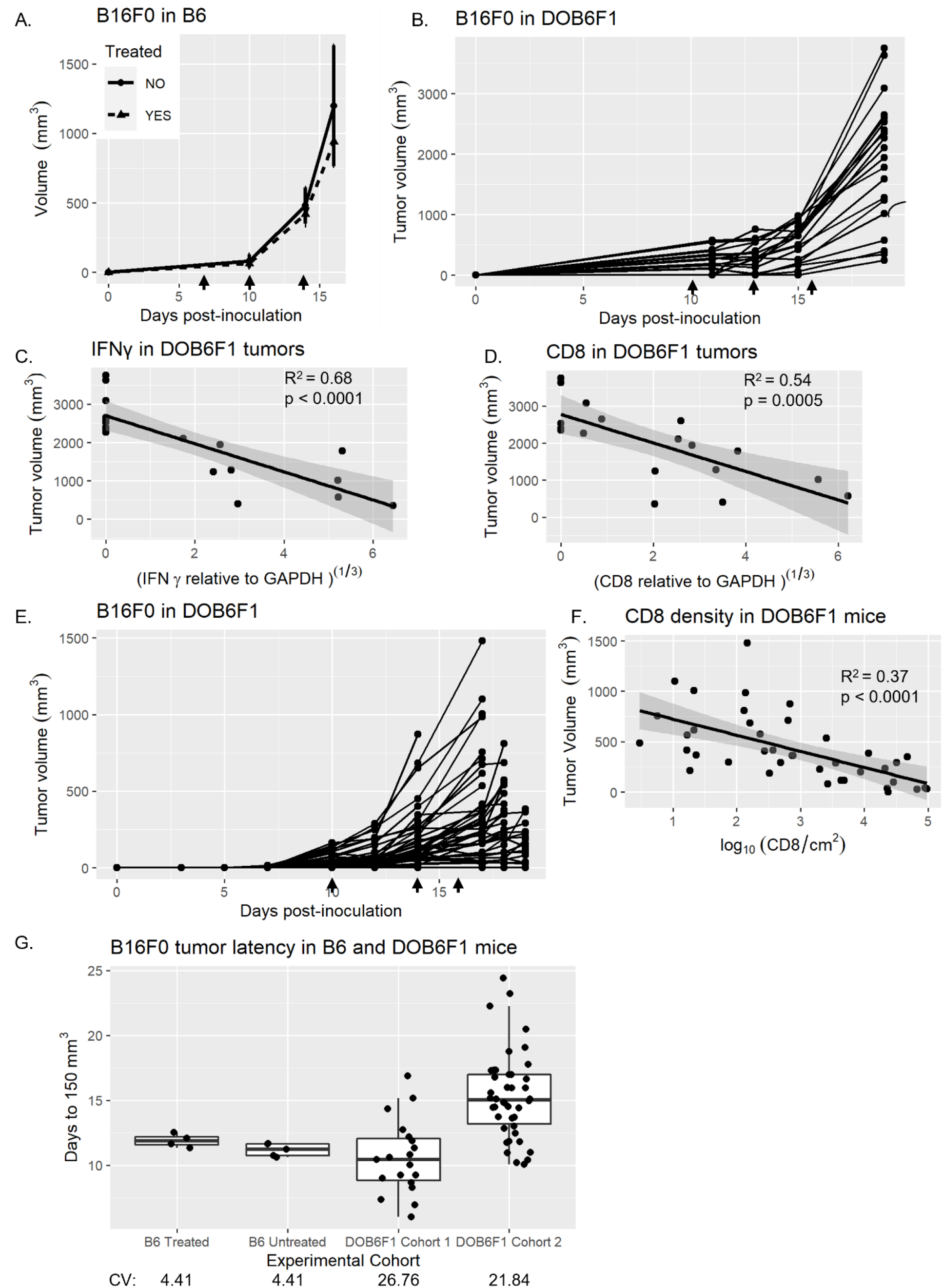
Host genome regulates response to combined ICI against B16F0 tumors. **A)** B16F0 tumor volume (mean mm^3^ ± SE) in B6 mice inoculated subcutaneously with 2×10 B16F0 cells on day 0 and treated with 200 μg anti-PD-1 and 100 μg anti-CTLA-4, on days 7, 10, 14 (n=4, dashed line) compared to untreated controls (n=5, solid line). **B)** Tumor growth in DOB6F1 mice (n=20) inoculated with 2×10 B16F0 cells and treatment with combined ICI on days 10, 13, and 16. Tumors harvested from **(B)** on day 19 were evaluated for abundance of **C)** IFNγ and **D)** CD8 transcripts via qRT-PCR relative to GAPDH, plotted against final tumor volume. Correlation was evaluated using Pearson’s correlation. **E)** Tumor growth in DOB6F1 mice (n=45) inoculated with 2×10 B16F0 cells and treatment with combined ICI on days 10, 13, and 16. **F)** Tumor infiltration of CD8, from **(E)**, measured by IHC plotted against final tumor volume. Correlation was evaluated using Pearson’s correlation. **(G)** Comparison of tumor latency (time to reach 150 mm^3^ in B6 and DOB6F1 mice bearing B16F0 tumor. Coefficient of variation (CV) of tumor latency is shown below each cohort.

**Table 1.** DO mouse cohorts used in study. Table showing number of DO dams, number of animals, and endpoint of cohorts used in study.

| Cohort | Treated n | Untreated n | DO Dams | ICI days | Endpoint |
| --- | --- | --- | --- | --- | --- |
| 1 | 20 | 0 | 12 | 10, 13, 16 | Post-ICI tumor immune analysis |
| 2 | 45 | 0 | 17 | 10, 13, 16 | Post-ICI tumor immune analysis |
| 3 | 96 | 19 | 31 | 3, 6, 10 | Efficacy, GWAS |
| 4 | 46 | 15 | 17 | 3, 6, 10 | Efficacy, GWAS |

To further test the role of host genetics in ICI outcomes and identify genetic loci associated with ICI response by linkage analysis, the study was scaled up with two separate biological replicate cohorts of DOB6F1 mice (Cohorts 3 and 4, **Table 1**) including both treated and untreated animals (total n=142 ICI treated, n=34 untreated). In an effort to increase response rates, treated mice received combined anti-CTLA-4/PD-1 ICI on days 3, 6, and 10 after subcutaneous B16F0 inoculation (**Fig. 2A**). Tumor latency for each mouse was then calculated by interpolating the day that the tumor reached 150 mm^3^ (a volume that can be reliably measured with calipers) from a linear regression of log-transformed tumor volumes. In treated mice response rates to ICI (**Fig. 2A**) were widely variable (latency range 8.7 - 65.9 days, where detectable tumors developed), compared to untreated mice where all tumors developed within a similar short time frame (latency range 6.19 - 21.13 days) (**Fig. 2B**). Notably, we observed 19 mice (~13%) from 14 different DO dams showing complete response (CR), as evidenced by no tumor development, and 43 mice (~30%) showing partial response (PR) defined as prolonged tumor latency versus controls (**Fig. 2C**).

**Figure 2.**
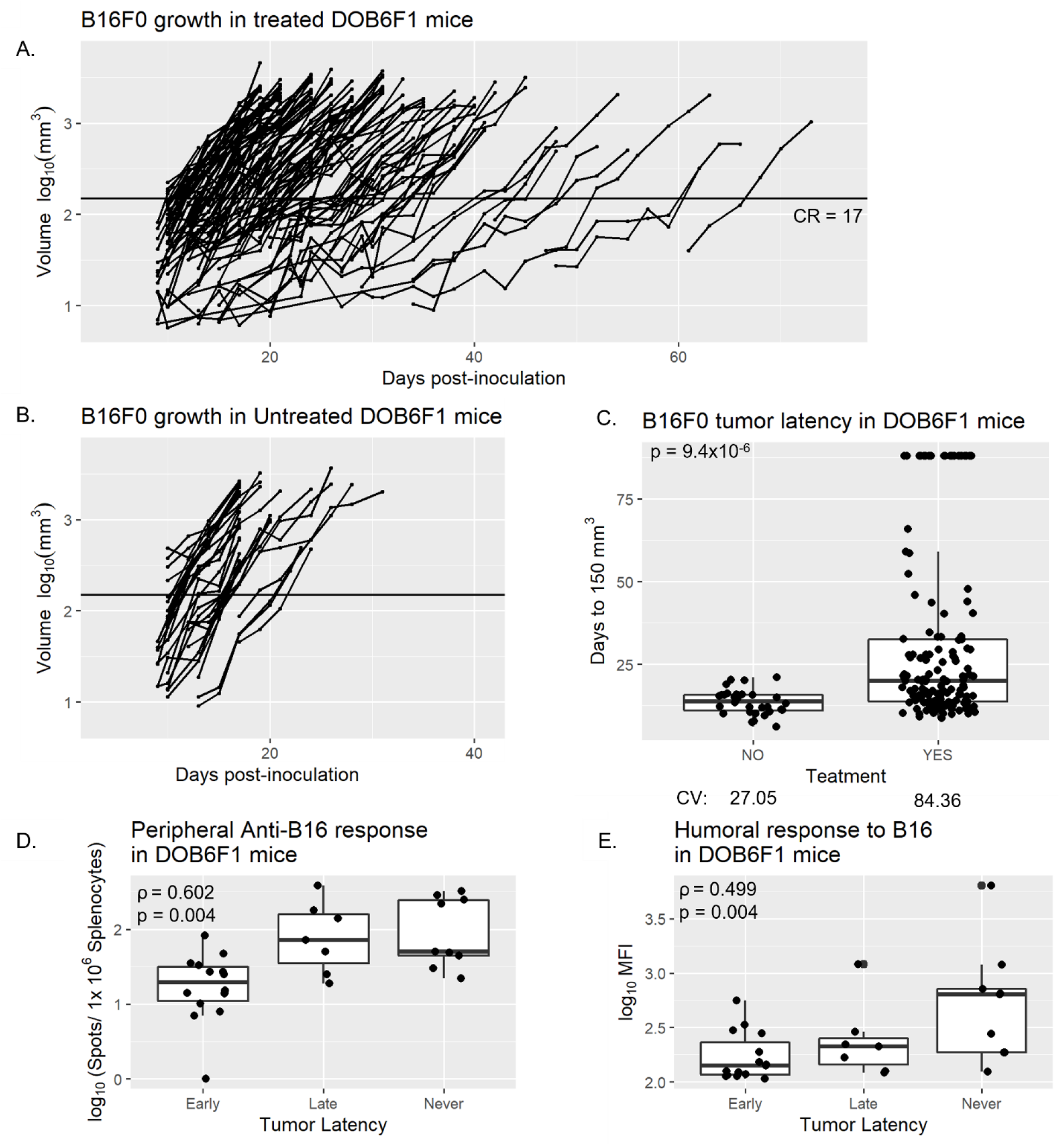
Response to ICI correlates to anti-tumor immune measurements in DOB6F1 mice. Log_10_ transformed tumor volume after subcutaneous inoculation with 2×10^5^ B16F0 cells in **(A)** ICI treated (200 μg anti-PD-1 and 100 μg anti-CTLA-4) and **(B)** untreated DOB6F1 mice. ICI treatment was administered on days 3, 6, and 10. Horizontal line indicates tumor volume of 150 mm^3^. **C)** Comparison of the tumor latency days to reach 150 mm^3^ in treated (n=142) and untreated (n=34) DOB6F1 mice, where tumors developed. Statistical significance determined via Student’s t-test. CV shown beneath each group. **D)** Anti-B16F0 T-cells were measured by IFNγ ELISPOT in treated DOB6F1 mice based on latency: early (<20 days), late (>40 days), or never developing tumor. Correlation was tested by Spearman’s rank correlation. **E)** Mean Fluorescence Intensity (MFI) of anti-B16F0 IgG detection in serum as measured by flow cytometry with grouping and correlation as described in **(D)**.

### Response to ICI in DOB6F1 mice is immune-mediated

All CR mice in Cohorts 3 and 4 (n=19) were given a contralateral re-challenge of B16F0 on day 88, three weeks after the final tumor developed, suggesting these mice had protection from primary tumor. All nineteen mice were protected from re-challenge, never forming a palpable tumor, supporting the presence of peripheral adaptive anti-tumor immune memory. In Cohort 4, serum from CR mice was collected prior to re-challenge to avoid artificial boosting of anti-tumor IgG. Adaptive anti-tumor T cells were evaluated in this cohort via IFNγ ELISPOT by testing splenocyte recall response to irradiated B16F0 cells. Production of B16F0-specific spots correlates to tumor latency, suggesting ICI response may involve adaptive anti-tumor T cell induction (**Fig. 2D**). Anti-tumor serum IgG levels followed a similar pattern, where likelihood of mounting a response increased in groups with either prolonged tumor latency or complete elimination (**Fig. 2E**). Interestingly, development of either anti-tumor IgG or anti-tumor T cells did not always occur in parallel (**Fig. S1C**),suggesting that multiple immune mechanisms contribute to ICI response in this genetically heterogeneous model. The intratumoral landscape of cohorts 3 and 4 was evaluated by IHC to compare the infiltration and distribution of cytotoxic CD8^+^ T cells and FoxP3^+^ regulatory T cells in a subset of early-forming (latency <20 days, n=14) and late-forming (latency >40 days, n=7) tumors. Surprisingly, unlike tumors collected within three days of the final treatment (**Fig. 1E, F**, Cohorts 1,2) there was no detectable correlation of either individual cell counts or ratio of CD8:FoxP3 to ICI outcome (not shown). We suspect this may be in part due to the timing of tissue collection, when tumor burden often exceeded 1,000 mm^3^ when immune infiltrates were scarce. Additionally, to identify pre-treatment biomarkers of ICI outcomes, leukocyte populations were evaluated from Peripheral Blood Mononuclear Cells (PBMCs) prior to tumor inoculation in Cohort 4. While the proportions of cell populations differed between DOB6F1 mice, we found no correlation between any of the measured markers relative to ICI treatment response (**Fig. S1B**).

We observe no difference in either tumor latency or survival when stratified by sex (**Fig. S2A**). None of the mice exhibited outward signs of autoimmunity or inflammatory side effects, such as colitis or dermatitis. The lack of autoimmune side effects may be attributed to the short treatment and survival window. We find no association of tumor latency with cage grouping, which may serve as an indirect measure of microbiome and other environmental influencers. There is an observable clustering within siblings, which speaks to the genetic regulation of this phenotype (**Fig. S2B**).

### Genetic linkage analysis reveals host genomic loci associated with ICI response

To perform a GWAS, we selected tumor latency as the phenotype representing response to ICI. As described above, tumor latency is defined by the time (in days) for a tumor to reach a volume of 150 mm^3^ (**Fig. 2A**). Genotyping was performed via GigaMUGA, and haplotype reconstruction and GWAS was performed using the R/qtl2 package (*29*, *30*). Genetic linkage analysis identifies peaks on Chr 2, 7, 13, and 14 with logarithm of odds (LOD) scores (an adjusted measurement of significance) greater than 6, with the strongest association in Chr13 (**Fig. 3A**). Importantly, we did not detect a correlation of the proportion of host B6 genome to tumor latency (**Fig. 3B**), suggesting variation in response does not result from “degree of foreignness” of the tumor. Similarly, the proportion of the other 7 contributing host founder strains also lacked correlation to tumor latency (**Fig. S3**).

**Figure 3.**
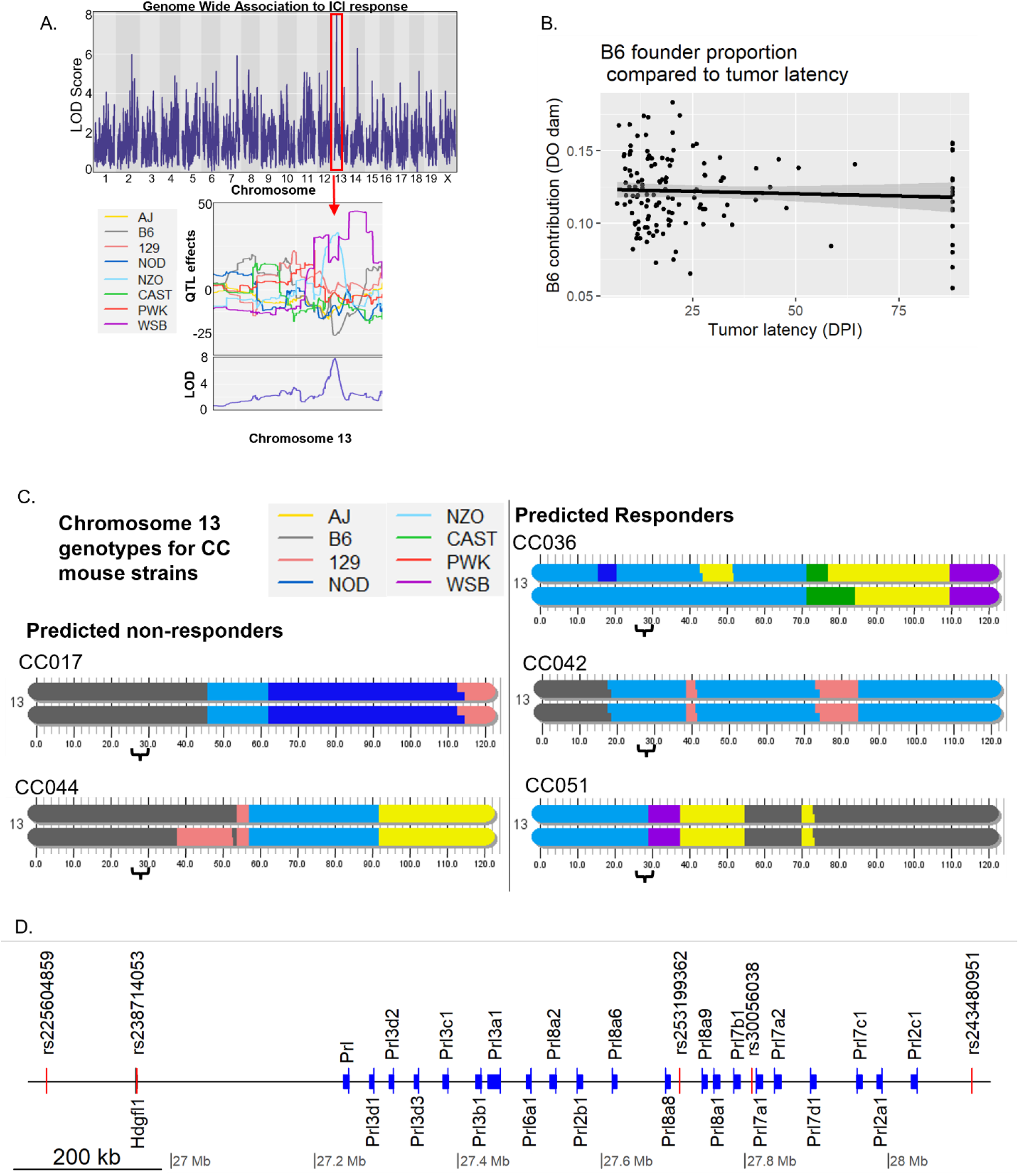
Genetic linkage analysis identifies genomic loci associated with response to ICI in B16F0-bearing mice. **A)** Manhattan plot for tumor latency **(Fig. 2A**) shows logarithm of odds (LOD) scores spanning the murine genome. Below, QTL effects of the strongest peak in Chr13 indicate the NZO genotype is a positive driver and B6 genotype is a negative driver of association. **B)** Plot of tumor latency against B6 contribution from DO dam. Correlation evaluated using Pearson’s correlation. **C)** Chr13 genotypes of CC mice selected for locus (indicated with brackets) validation. Modified from (13). **D)** The 5 highest associating polymorphisms (red) plotted against murine prolactin genes (blue) and Hdfgl1 gene (black) in the Chr13 locus. Gene start sites are marked by a thin vertical line with bar extending in gene direction.

### Validation of the prolactin locus as a regulator of ICI response

Within the Chr13 peak, analysis of QTL effects by founder revealed that the association is driven positively by the NZO genotype and negatively by the B6 genotype (**Fig. 3A**). The negative association with B6 is consistent with the lack of ICI response to B16F0 in inbred B6 mice (**Fig. 1A**). Whole-genome analysis, derived from haplotype reconstruction data, shows the SNPs with highest LOD scores are located within the genomic region containing the mouse prolactin gene family (**Fig. 3D**). All 5 of the SNPs identified are specific to the NZO genotype and are not shared by any of the other founders (**Fig. 3D**).

To validate the locus on Chr13 we identified CC mouse strains that are either putative non-responders to ICI, containing the B6 genotype at this locus (CC017 and CC044), or putative ICI responders with the NZO genotype in this region (CC036, CC042, and CC051). These strains were selected to also lack confounding driver genotypes within any of the other 3 identified strong loci on chromosome 2, 7, and 14 (**Fig. 3A**). CCB6F1 mice were inoculated with B16F0 and treated with ICI as described above. CC044B6F1 and CC017B6F1 showed no response to ICI, as predicted based on genotype, whereas ICI-treated CC042B6F1 and CC051B6F1 had reduced tumor burden at day 15 (p=0.002, p < 0.001, respectively) (**Fig. 4A**). Despite being a predicted responder, CC036B6F1 mice fail to respond to ICI (**Fig. 4A**). A similar trend was observed in tumor latency measurements **(Fig 4B)**. This is consistent with the individual DOB6F1 genotypes, where ~24% of mice identified as NZO within this locus were characterized as non-responders with tumor latency matching untreated mice. Of note, ~47% of DOB6F1 mice that were the NZO genotype within the Chr13 locus had a CR or late forming tumor (tumor latency >40 days). The fact that the NZO genotype in this locus lacks complete penetrance perhaps speaks to the multigenic nature of anti-tumor immune response with ICI.

**Figure 4.**
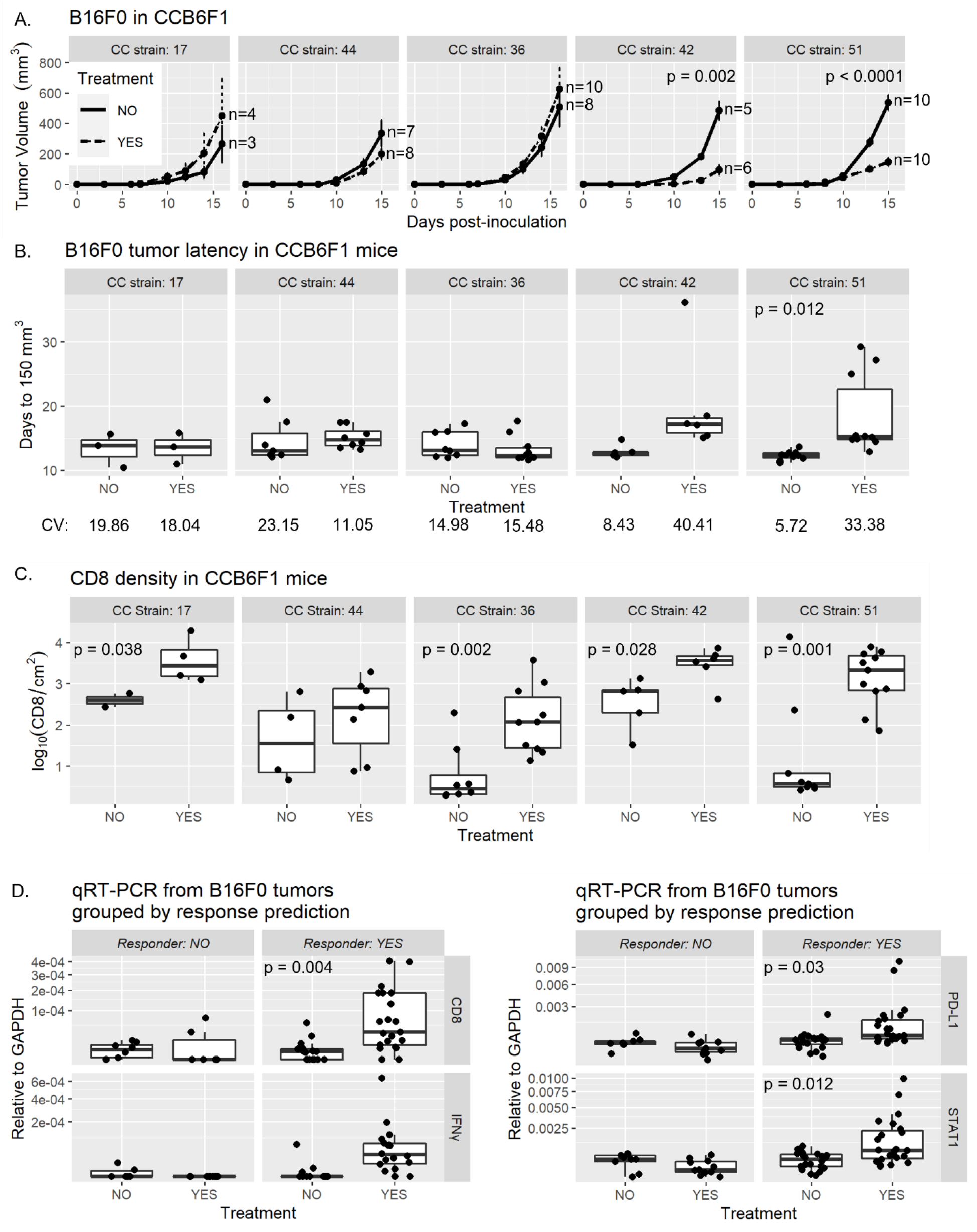
Validation of the Chr13 locus in CC mice. **A)** Tumor volumes (mean mm^3^ ± SE) of CCB6F1 with (dashed lines) and without (solid lines) ICI treatment on days 3, 6, and 10. Graphs show group averages up to the first euthanasia event. Significance determined by Student’s T-test of tumor volume at day 15 or 16. **B)** Comparison of the tumor latency, days to reach 150 mm3, in treated and untreated CCB6F1 mice grouped by CC strain. Statistical significance determined via Student’s t-test. CV shown beneath each group. **C)** Tumor infiltration of CD8, from **(A)**, measured by IHC grouped by CC strain and treatment. Significance determined by Student’s T-test. **D)** Averaged qRT-PCR of CD8, IFNγ, and downstream IFNγ response genes PD-L1 and STAT1 results from grouped predicted non-responder mice (CC44, CC17) versus predicted responder mice (CC36, CC42, CC51). Significance determined by Student’s T-test.

To investigate whether the ICI impact on tumor burden was immune-mediated, we utilized IHC to determine CD8 infiltration. We found that all three predicted responder strains show significant increases in CD8 infiltration following treatment **(Fig 4C).** We also observed a significant increase of CD8 infiltrates within the non-responder CC017. However, qRT-PCR from these tumors have no detectable IFNγ expression in either the untreated or treated mice **(Fig. S4A)**. We detected a consistently higher abundance of anti-tumor immune-related transcripts including CD8 and IFNγ, and IFNγ-driven transcripts PD-L1, and STAT1 within treated versus untreated control mice from the responding strains **(Fig. S4A)**. In individual strains, significant differences were found within CC051 and CC044, with CC051 showing significantly elevated CD8 and STAT1 transcripts after treatment and surprisingly CC044 having significantly lowered CD8 and STAT1. When predicted responder and non-responder strains are grouped together, CD8, STAT1, and PD-L1 transcripts are significantly higher after treatment (**Fig. 4D**). Taken together, these results support the hypothesis that the NZO genotype within the prolactin region of Chr13 influences ICI-induced immune response. Anti-tumor antibodies were also measured using flow cytometry as described above, and only CC051 showed a significant induction of anti-B16 antibodies **(Fig. S4B)**.

During the locus validation experiments we selected the CC007 strain. We found this predicted non-responder showed an unexpected positive response to ICI, demonstrated by increased tumor latency with 1/6 mice failing to develop a tumor until ~60 days (**Fig. 5A**). A potential explanation of this finding stems from a secondary QTL analysis performed as a binary response variable using either presence or complete absence of tumor development, instead of the continuous variable of tumor latency. This was done post-hoc to search for potential confounding loci after CC strains 036 and 007 did not respond as predicted. In this context, we find de-emphasis of some peaks from the continuous variable analysis, with emergence of several novel peaks including a strong peak on Chr6 (**Fig. 5B**, upper panel). Importantly, in the binary analysis the peak and QTL effects from the Chr13 locus are still maintained. The novel peak within Chr6 shows the CAST or NOD genotypes are negative drivers of the effect and A/J is a positive driver (**Fig. 5B**, lower panel). CC007 has the B6 genotype in Chr 13, suggesting a non-responder, but also has the A/J genotype in the Chr6 loci confounding the prediction **(Fig 5C)**. To further explore the Chr6 locus we selected CC008, which also has the A/J genotype in the Chr6 locus, with no other known confounding locus genotypes **(Fig 5C)**. In CC008B6F1 mice receiving ICI, we observe a tumor onset pattern similar to CC007B6F1, with most mice showing minimal response to therapy, but 2/7 achieving complete response. We also found a significant increase of CD8^+^ tumor infiltrates after ICI in CC007B6F1 mice by IHC (**Fig. 4E**). This trend is also present in tumors from CC008B6F1 mice, but is not significant, potentially due to the lack of tissue from mice that fully eliminated the tumor. We also observed similar trends of increased inflammatory markers detected via qRT-PCR from flash-frozen tumor sections **(Fig. S4A**). These findings warrant further exploration of the Chr6 locus and its influence on complete response to ICI therapy. Additionally, these results stress the importance of phenotype selection for GWAS, and further demonstrate the multigenic nature of anti-tumor immunity.

**Figure 5.**
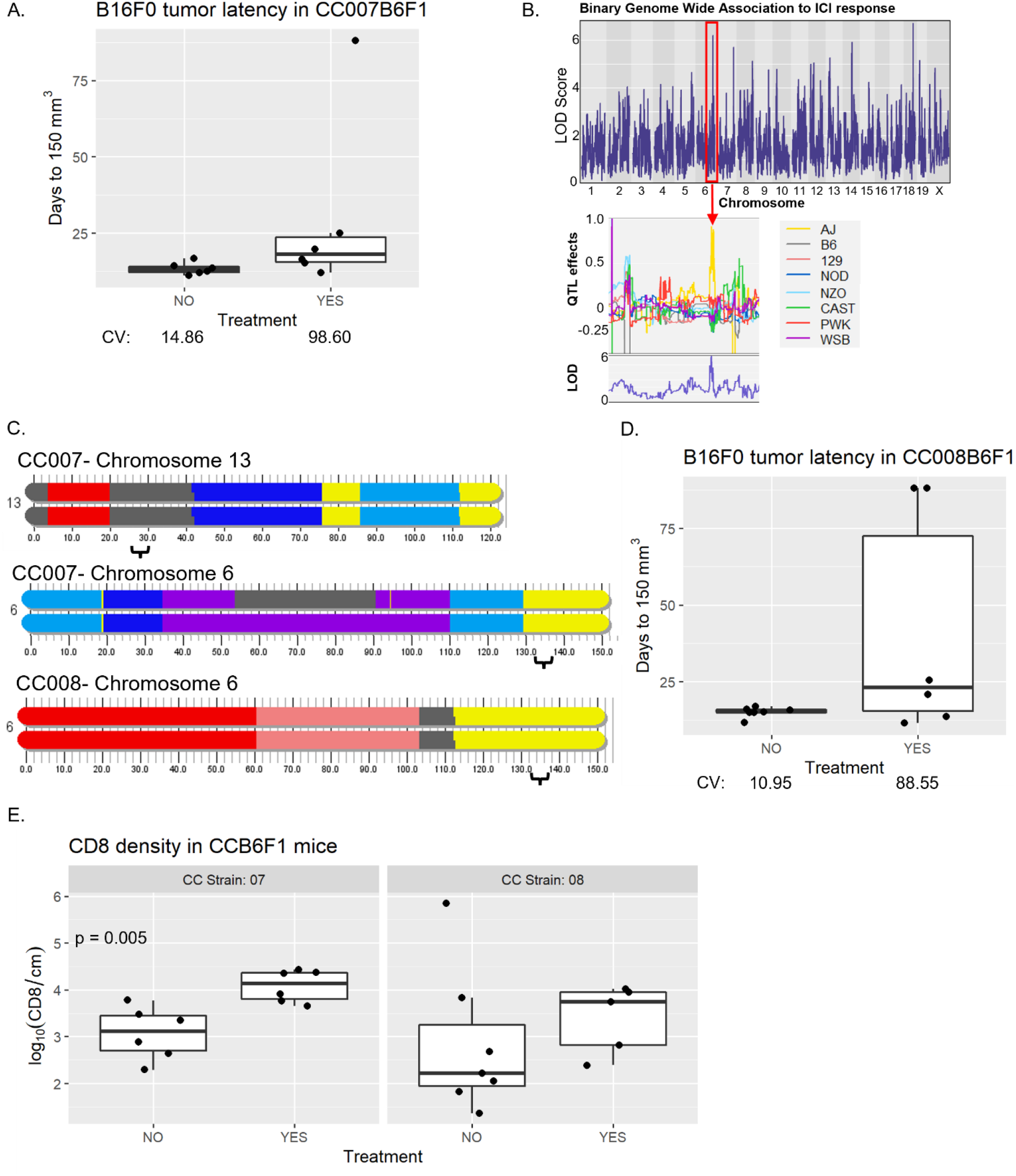
Binary genetic linkage analysis reveals additional genomic loci associated with ICI response against B16F0. **A)** Comparison of the tumor latency, days to reach 150 mm^3^, in treated and untreated CC007B6F1 mice. CV shown beneath each group **B)** Manhattan plot from binary analysis of presence or absence of tumor development in treated DOB6F1 mice **(Fig. 2A)**. QTL effects for the Chr6 peak indicate that the A/J genotype is a positive driver for complete response. **C)** Chr6 genotype in CC007 and CC008 mice (bracket indicates locus of interest). Chr 13 genotypes also provided for CC007. Modified from (13). **D)** Comparison of tumor latency, days to reach 150 mm^3^, in treated and untreated CC008B6F1 mice. CV shown beneath each group. **E)** Tumor infiltration of CD8, from **(A/D)**, measured by IHC grouped by CC strain and treatment. Significance determined by Student’s t-test.

### Prolactin co-administration augments ICI response

To directly investigate whether prolactin has an immunomodulatory function in the context of ICI treatment, we combined ICI with mild hyperprolactinemia in inbred B6 mice, which historically fail to respond to combined ICI alone ((*28*) and **Fig. 1A**). Systemic prolactin levels were elevated with a subcutaneous 30-day slow-release prolactin pellet as described previously (*31*), with placebo matrix-only pellets as controls. Seven days after pellet implantation, allowing time for equilibrium of prolactin release, mice were inoculated with B16F0 and treated with combined ICI on days 3, 6, and 10 (**Fig 6A**). Compared to treated mice receiving the placebo pellet, mice receiving prolactin with ICI showed an interaction effect, with lower tumor burden in 5/8 dual-treated mice by day 19 after inoculation (**Fig. 6B**). This was demonstrated in a linear mixed effects model analyzing growth trajectories on log-transformed tumor volume. To account for different growth trajectories over the course of the study, a random slope and intercept were estimated for each animal while day, treatment and pellet type were analyzed as fixed effects. Interactions between the fixed effects were also examined for statistical significance. From this analysis, we see a borderline significant interaction effect (p=0.0684) between ICI treatment and pellet type, indicating that ICI treatment influenced tumor growth in the prolactin group versus the placebo group. The significance of these results are lessened by one mouse in the ICI with placebo pellet group with abnormally low tumor burden compared to our historical controls in inbred B6 mice (**Fig. 1A**). If we exclude this outlier, the interaction effect of ICI treatment in the prolactin pellet group is much stronger (p=0.0032). Of note, in the absence of ICI we also detect a small, insignificant increase in tumor volume in mice receiving prolactin versus placebo pellet (1438 ± 934 mm^3^ and 818 ± 282 mm^3^, respectively, at day 19). The trend is also visible in tumor latency measurements (**Fig 6C**).

**Figure 6.**
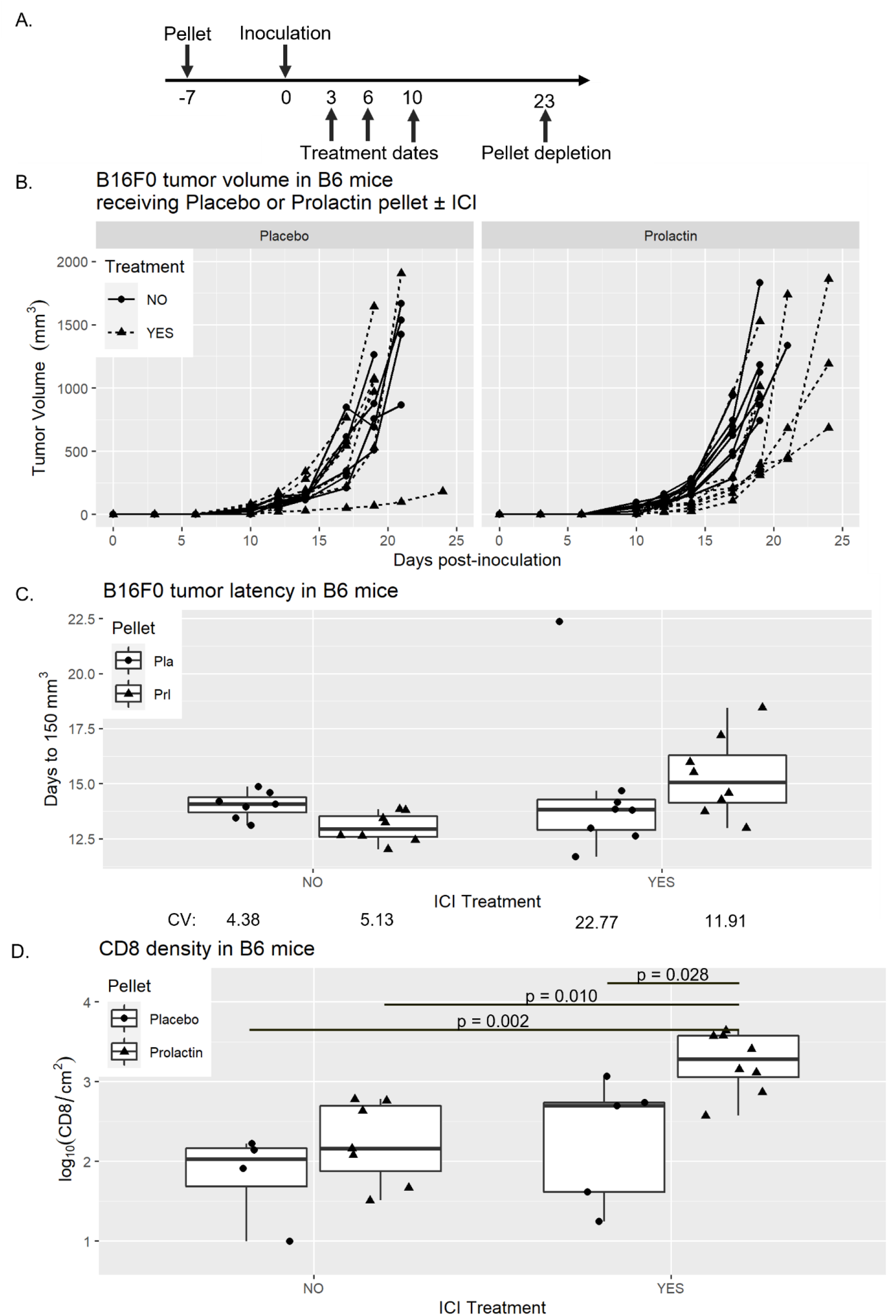
Systemic prolactin elevation augments combined ICI therapy in B16F0-bearing B6 mice. **A)** Schematic showing dates of pellet implantation, treatment dates, and approximate pellet depletion. All dates relative to tumor inoculation at day 0. **B)** B16F0 tumor volume in B6 mice receiving a subcutaneous placebo pellet (left) or 30-day slow-release prolactin pellet (right) 7 days prior to inoculation. Mice were treated with combined ICI on days 3, 6, and 10 after tumor inoculation **C)** Comparison of the tumor latency, days to reach 150 mm^3^, in treated and untreated B6 mice with shape indicating pellet implant. CV shown beneath each group **D)** Tumor infiltration of CD8, from **(B)**, measured by IHC. Significance determined by Tukey’s HSD post-hoc test following Two-way Anova with interaction for ICI treatment and pellet implantation.

To determine if there were changes in the tumor microenvironment we examined collected tumors for CD8 infiltrates via IHC and observed that mice receiving the prolactin pellet and ICI had significantly higher CD8 infiltration than any other group (**Fig 6D**). We also performed qRT-PCR on collected tumor sections. We see a non-significant trend of increasing expression of CD8 from mice receiving no therapy with similar trends in IFNγ and downstream IFNγ response genes PD-L1 and STAT1 (**Fig. S5**). This suggests that combined prolactin with ICI is inducing an inflammatory milieu within the tumor, supporting the observed interaction effect. We also examined peripheral T cell response to B16 using ELISPOT in spleens from mice receiving ICI alone or prolactin and ICI, however no induction of peripheral T cells was detected (not shown). There was also a lack of induction of anti-B16 serum IgG measured by flow cytometry (not shown). The lack of detectable peripheral anti-B16 immunity may be the result of the short time frame of the experiment, the persistence of the primary tumor, or additional immune tolerance mechanisms in B6 mice. Taken together, these data show that systemic prolactin augments ICI efficacy and enhances the inflammatory environment within B16 melanoma.

## DISCUSSION

ICI has demonstrated clinical success, particularly in the settings of metastatic melanoma and lung cancer. These primarily carcinogen-driven malignancies, which result in high tumor mutational burden and subsequent neoantigen presence (*16*), should theoretically produce high response rates to ICI, however many patients fail to benefit. Multiple studies have focused on the tumor and the tumor microenvironment to explain this divergence (reviewed in (*32*)). In this study, we focused on the regulation of the host immune response, leaning on decades of autoimmunity research and the numerous links between genetics and susceptibility to autoimmune conditions (reviewed in (*33*)). Evaluation of host genome and ICI outcomes is thus far almost exclusively limited to examination of HLA, where HLA diversity and specific HLA subtypes have been associated with ICI outcomes (*34*, *35*). In our current study, we utilized the DO system to reduce extraneous variables and evaluate the entire host genome in a less-biased manner, elucidating host-intrinsic pathways that influence response to ICI. Interestingly, there is no obvious peak within the mouse MHC locus on Chr17, suggesting MHC or MHC-peptide combinations may not be the strongest influencer of ICI response in this model. This approach revealed that systemic prolactin treatment results in augmentation of ICI response against the B16F0 melanoma model. This result, supported by previously published data associating prolactin polymorphisms with human autoimmune diseases, justifies further investigation into prolactin signaling as a method to augment ICI. The exact timing, location, and cell populations involved should be explored. Further, an evaluation of whether prolactin promoter SNPs in the human population correlate to ICI outcomes should also be investigated.

Prolactin may have differential effects on tumor cells as compared to immune cell populations. Outside of our data and others showing the stimulatory effect prolactin can have on immune cells, there is evidence that prolactin has tumorigenic and pro-tumor growth factor activities. Described initially in breast cancer, prolactin may enhance both growth and metastasis (reviewed in (*36*)). Additional research has shown prolactin signaling may play a role in prostate, ovarian, pancreatic, and endometrial cancers (*37–40*). In our data, mice that received the prolactin pellet alone show a slightly higher average tumor burden by day 19 than mice receiving the placebo pellet without ICI. Future studies of prolactin signaling as an anti-tumor immune stimulus must also consider the potential counteractive role prolactin can have on cancer growth.

Prolactin alone was insufficient to eliminate tumor with ICI in inbred B6 mice. This could be due to heightened self-tolerance observed in the B6 strain. Prolactin alone was inadequate to induce a SLE like phenotype in B6 compared to BALB/c mice. In B6 mice they needed an additional genetic susceptibility to break tolerance. Higher self-tolerance in B6 mice is also supported by tumor immunogenicity study which shows that tumors in B6 were less immunogenic than tumors in BALB/c mice (*26*).

There are several advantages to modeling tumor immunotherapy in DOF1 crosses versus the human population. DOF1 mice have reduced environmental confounders - all mice are of similar age, have controlled living conditions, and are on the same diet. All subjects receive an identical treatment program and, critically, a genetically identical tumor line, which should control for tumor heterogeneity. This is underscored by comparison of the tumor latency Coefficient of Variation (CV) in the inbred B6 mice versus DOB6F1 mice. Inbred B6 mice treated with ICI on days 3,6,10 (**Fig 6B**) have a CV of 22.77 compared to DOB6F1 mice on the same treatment schedule (**Fig 2C)**, which have a CV 84.36. These CVs are significantly different, as determined by modified signed-likelihood ratio test using the R cvequality package (p = 0.002) (*41*, *42*). Among CCB6F1 strains, we found high tumor onset CVs in CC051, CC007, and CC008. However, the degree of ICI response in these mice appears to follow a binary rather than continuous trend. This may indicate other genetic features, such as residual heterozygosity, which was shown to affect onset of colitis in CC mice (*43*). The CC mice we selected do not have known heterozygosity in the loci with the highest LOD scores, but other minor loci may contribute. Additional environmental, hormonal, or other non-genetic influencers may also be involved in observed phenotypes.

Utilization of a DOF1 cross versus the DO stock reduces the degrees of freedom for genotype predictions from 36 to 8. This can reduce the number of animals needed for a study, but also obfuscate recessive traits. This model may easily be extended to investigate other aspects of tumor immunology, including immune surveillance or alternative immunotherapy modalities. Our validation studies in CCB6F1 mice have also revealed new models with a range of responses to ICI against genetically identical tumors (**Fig. 4**), which may be advantageous versus comparing multiple tumor models in multiple inbred mouse strains, as has become common practice (*26*). These and other CCF1 models may bolster future studies of regulatory mechanisms of anti-tumor immune activity. Recent work from others have also highlighted the utility of the DO stock and CC lines in models of infection, vaccination, autoimmunity, and cancer susceptibility (*44–50*).

Over the course of the study, we saw no evidence that cage grouping or sex influences response to ICI (**Fig. S2A**). Cage grouping has a significant effect on microbiome composition and the lack of clustering of responders within cages suggests that differences in the microbiome did not influence outcomes in our study (*51*). The gut microbiome is an area of intense investigation for ICI response and may reveal predictive biomarkers and therapeutic strategies to improve ICI success rates including microbiota transplants (*52*). Future investigation using DOF1 or CCF1 mice could provide an opportunity to investigate microbiome influence on ICI, and yield insight into interactions between the genome, the microbiome, and anti-tumor immunity.

Despite the reduced number of test subjects needed for GWAS studies in DO mice versus humans, DO mouse studies may still require large numbers of animals depending on the phenotype chosen for analysis. Minor allele frequency in the DO/CC mouse populations is a factor that reduces the required “n” because the minor allele frequency is close to 0.125 compared to the large number of near zero minor allele frequencies in humans (*53*). Ideal candidate phenotypes will have low variation within a particular inbred model, with variation between different inbred strains. Importantly, not all phenotypes will have a genetic influence (*54*, *55*). The effect size of the variation in phenotype is also an important consideration when determining sample size. Additionally, while genetics may regulate a specific phenotype in mice, the genetic component itself may not translate to humans. However, identified genes and their related pathways may still impact the same biological processes, and thus modulation of these targets may still have clinical value.

Complex traits are also multigenic traits. This is evident in our study, where multiple genomic loci were identified, and the driver genotypes for our most prominent locus do not uniformly predict outcomes. Additionally, we find variation in intratumoral immune profiles and peripheral anti-tumor T cell and humoral responses. Any large GWAS study has the potential to reveal multiple loci that contribute to an observed phenotype, and thus validation of suspected loci should account for this. In this study, we utilized multiple CC mouse strains containing the driver genotype at one specific locus, while avoiding strains with drivers at other prominent loci. This effect is especially highlighted in our results with CC036B6F1, a predicted responder that failed to respond, and CC007B6F1, which contains the negative driver genotype at the Chr13 locus, but a positive driver in Chr6 discovered when tumor development was tested as a binary trait. The failure of strain CC036B6F1 to respond despite significant T-cell infiltration in treated tumors may be due to other genetic loci controlling the immune response. In a study investigating cancer susceptibility in CC mice, 100% of CC036 mice developed spontaneous tumors and one of the associated loci contained the Nfkb1 gene (*50*). Combined with our study, these results suggest the CC036 strain may exhibit immune deficiencies, particularly for cancer surveillance. It may not be possible to completely mitigate all contributing loci, which is why we chose multiple CC strains. Additionally, the number of CC strains needed to validate a locus may be dependent on the strength of the association, and this should be considered when planning validation studies.

In this study we have pioneered the use of F1 crosses of DO and CC mice for use with strain-specific syngeneic tumors to further our understanding of tumor immunology. The DO mouse population serves as a valuable surrogate for the human population showing a wide variation in treatment effectiveness, and unlike the human population, it is easy to evaluate genetic loci associating with response. The utilization of CC mouse lines identified in the genetic screen serves a dual purpose for testing loci for phenotype influence and generation of new models that can replicate therapeutic outcomes. We utilized the DO and CC models, which identified a locus within the mouse prolactin family that associated with ICI response after challenge with B16F0 melanoma. Furthermore, our data shows that using this approach, we developed CC models that can be used to replicate response and non-response to therapy which can serve in future investigations.

## MATERIALS AND METHODS

### Mice and Cell lines

All animal handling and experiments were conducted in accordance with guidelines and regulations set by the Wayne State University Institutional Animal Care and Use Committee (IACUC). Both male and female mice were utilized in this study. B6 and DO mice were purchased from the Jackson Laboratory, all CC mice were purchased from theSystems Genetics Core Facility at the University of North Carolina (*13*). CC mouse lines were generated and bred at Tel Aviv University in Israel (*56*), Geniad in Australia (*57*), and Oak Ridge National laboratory (*58*) prior to relocation to UNC. For the CC lines used in this study, the full nomenclature is as follows: CC007/Unc, CC008/GeniUnc, CC017/Unc, CC036/Unc, CC042/GeniUnc, CC044/Unc, CC051/TauUnc. On average, we selected 2-3 DOB6F1 siblings per DO dam in the treated cohorts, and 1 for the untreated cohorts, to increase genetic diversity. B16F0 (ATCC CRL-6322™, Virginia USA) cells were cultures in a sterile environment at 37 °C and 5% CO_2_ using DMEM (Life Technologies, Grand Island NY) containing 10% FBS (R&D Systems, Flowery Branch, GA) and penicillin (100 U/mL), and streptomycin (100 μg/mL). Cells used in experiments were directly purchased from ATCC and maintained for fewer than 5 passages, and thus authentication is not necessary. All tumor growth experiments were performed on mice 6-8 weeks old with inoculations performed on the right inguinal region with 2×10^5^ B16F0 cells suspended in DMEM. Tumor growth was monitored by caliper measurement and tumor volume calculated as (W× W × L)/2(W× W × L)/2.

### Treatment with ICI

Mice were randomized into treated and untreated groups. ICI was given by intraperitoneal injection with 200 μg anti PD-1 (clone RMP1-14, Leinco Technologies) and 100 μg anti CTLA-4 (clone 9D9, Leinco Technologies) in volume of 200 μg 1X PBS.

### Blood collection

PBMCs were collected from retro-orbital bleeds using heparinized capillary tubes into 1.5 mL conical tubes containing 5 U heparin. Red Blood Cell lysis was performed via water lysis and remaining cells were immediately used. Blood for serum was collected without heparin and allowed to clot for 30 minutes before centrifugation 400 x g for 5 minutes to pellet cells and clotting factors.

### Induction of hyperprolactinemia

Ovine prolactin was obtained from the National Hormone & Peptide Program (NHPP) and was converted into a slow release pellet for subcutaneous implantation by Innovative Research of America to release 100 μg/day for 30 days to result in a mild hyperprolactinemia as described previously (*25*).

### Quantitative Reverse Transcription PCR

Tumor tissue from mice was flash-frozen in liquid nitrogen and homogenized in Trizol as described by manufacturer (Thermo Fisher, Waltham, MA). cDNA was synthesized using a ProtoScript First Strand cDNA synthesis kit (New England Biolabs, MA, E6300S) using PolyDT primers. qRT-PCR was conducted with Taqman probes (Thermo Fisher) for GAPDH (Mm99999915_g1), CD8α (Mm01188922_m1), IFN-γ (Mm01168134_m1), PD-L1 (Mm03048248_m1), STAT1 (Mm01257286_m1). 10 ng of cDNA/well was used and relative mRNA was calculated (2^-ΔCT^) relative to GAPDH. Transcripts that failed to amplify in all technical replicates were set to a CT value of 55. Transcripts in which only 1/3 of technical replicates amplified were removed. Statistical comparisons between groups was performed using Student’s T-test, with Welch’s correction where appropriate, where stated in figures.

### B16F0 specific Ab and T cell response

B16 reactive T cells were assayed by IFN-γ ELISPOT assay as previously described (*52*). ELISPOT plates were coated with anti-mouse IFN-γ antibody (clone AN18, Invitrogen) and incubated overnight at 4 °C. Plates blocked with DMEM with 10% FBS and 1% P/S for 2 hours at 37 °C prior to additions of 2×10^5^ splenocytes and 5×10^4^ irradiated B16F0 cells. Plates were incubated for 48 hrs at 37 °C and 5% CO2, before being washed and coated with biotinylated anti-mouse IFN-γ (clone R4-6A2, eBioscience™). Following detection plates were washed and then spots developed by AEC substrate. Dried plates were read using ImmunoSpot^®^ and counted using ImmunoSpot^®^ software. Anti-B16F0 IgG was measured by incubating serum from terminally bled mice and incubated with B16F0 cells. Detection was performed using PE anti-mouse IgG and MFI as primary readout.

### Flow cytometry of PBMCs

PBMC populations from DOB6F1 mice were performed on a BD LSRII flow cytometer (Becton Dickinson, Franklin Lakes, NJ). The following panel was used: Viability (Ghost Dye™ Violet 510, TONBO biosciences, CD3ε APC780 (clone 17A2), CD19 BV 605 (clone 6D5), CD4 AF 488 (clone RM4-5), CD8 BB700 (clone 53-6.7), FoxP3 eFluor 660 (clone FJK-16s). PBLs samples were gated based on viable singlets. B-cells were CD19^+^/CD3^-^, CD3^+^ defined pan T cells, CD4 T cells (CD3^+^/CD19^-^/CD4^+^), CD8 (CD3^+^/CD19^-^/CD8^+^), Double negative T cells (CD3^+^/CD4^-^/ CD8^-^), Regulatory T cells (CD3^+^/CD19^-^/CD4^+^/FoxP3^+^). Example gating can be found in **Fig. S7**

### Mouse genotyping and QTL analysis

Genotyping was performed using GigaMUGA(*29*) performed by Neogen (https://www.neogen.com/categories/custom-genomic-solutions/gigamuga/). QTL analysis was performed using the R/qtl2 (*30*) (v0.24) in R version 4.0.3. Briefly DOB6F1 Haplotypes were constructed using a hidden Markov Model and linkage mapping was performed using a mixed effects linear model with Sex and relatedness (as determined via kinship matrix) as co-factors. The binary analysis was performed in continuous mode since the binary mode cannot utilize the kinship matrix. Suggestive peaks were more closely examined for strain driven QTL effects using single QTL with additive allele effects. Single strain QTL effects were examined at suggestive peaks (LOD > 6) to determine putative positive and negative drivers of the association. CC mouse genomes were queried ± 1 Mb of suggestive peaks to determine dominant strain genotype and used to select CC mouse strains that were only a putative positive or negative driver at one peak. CC mouse genomes, status, and ideograms provided and maintained by UNC Computational Systems Biology (*59*, *60*). Identified SNPs were plotted with genes using R/genoplot (*61*).

### IHC and Weka Segmentation

CD8 infiltration was determined via Immunohistochemistry. Briefly Formalin Fixed Paraffin Embedded sections were deparaffinized and rehydrated in Xylene and Ethanol rinses prior to antigen decloaking via heat pressure for 10 sec 105 °C and 10 sec 95 °C. Normal horse serum was used for non-specific blocking and Rabbit anti-mouse CD8α (clone D4W27, Cell Signaling Technology^®^) was used as detection antibody. ImmPRESS^®^ Horse Anti-Rabbit IgG Polymer Kit (Vector Laboratories) was used for secondary antibody and chromagen reaction using the ImmPACT^®^ Vector^®^ Red kit (Vector Laboratories). Tissues were counterstained with Mayer’s Hematoxylin (Thermo scientific) and then dehydrated in Ethanol and Xylene rinses before being mounted with Permount^®^ (Fisher). To determine CD8 infiltration in tumor sample after staining slides were scanned with a SCN 400 Slide Scanner (Leica). Representative scan images were used to train a Weka

Segmentation (*62*) classifier on tumor, melanin, and positive staining. After segmentation images were converted to 8 bit and positive staining was set to red using image threshold. Images were then converted to black and white filtering out all features other than positive staining. Positive staining regions were filled in using a fill holes function and clustered objects were split using a watershed function. Positive cells were finally counted using a minimum circularity of 0.3 and a size exclusion of 200-20,000 pixels. A representative image showing the process from slide image to counted positive cells can be seen in **Fig S6**

## Acknowledgements

We thank Dr. Amalraj Thangasamy for his assistance with scanning microscopy slides. Dr. Karl Broman for his assistance with using R/Qtl2 and maintaining an active community around the software he developed. Drs Karin List and Jeffrey Frelinger for reviewing early versions of this manuscript and providing insight and suggestions. We also would like to thank the WSU Grid Team for their help in utilizing the High Performance Computing Grid.

## Funding

National Institutes of Health Ruth L. Kirschstein National Research Service Award T32-CA009531 (JBH)

National Institutes of Health R37 CA220482 (HMG)

National Institutes of Health P30 CA022453

DMC Foundation (HMG)

Pardee Foundation (HMG)

## Author contributions

Conceptualization: HMG, JBH

Methodology: JBH, HMG, JEG

Investigation: JBH, HMG, MCM, JEG, NM, JM, CM, MB, AF Formal Analysis: JBH, GD

Visualization: JBH, GD

Funding Acquisition: HMG

Project administration: HMG, MCM

Supervision: HMG, MCM

Writing – Original Draft: JBH, HMG

Writing – Review and Editing: JBH, HMG, JEG, MCM

## Supplementary Materials

**Fig S1.**
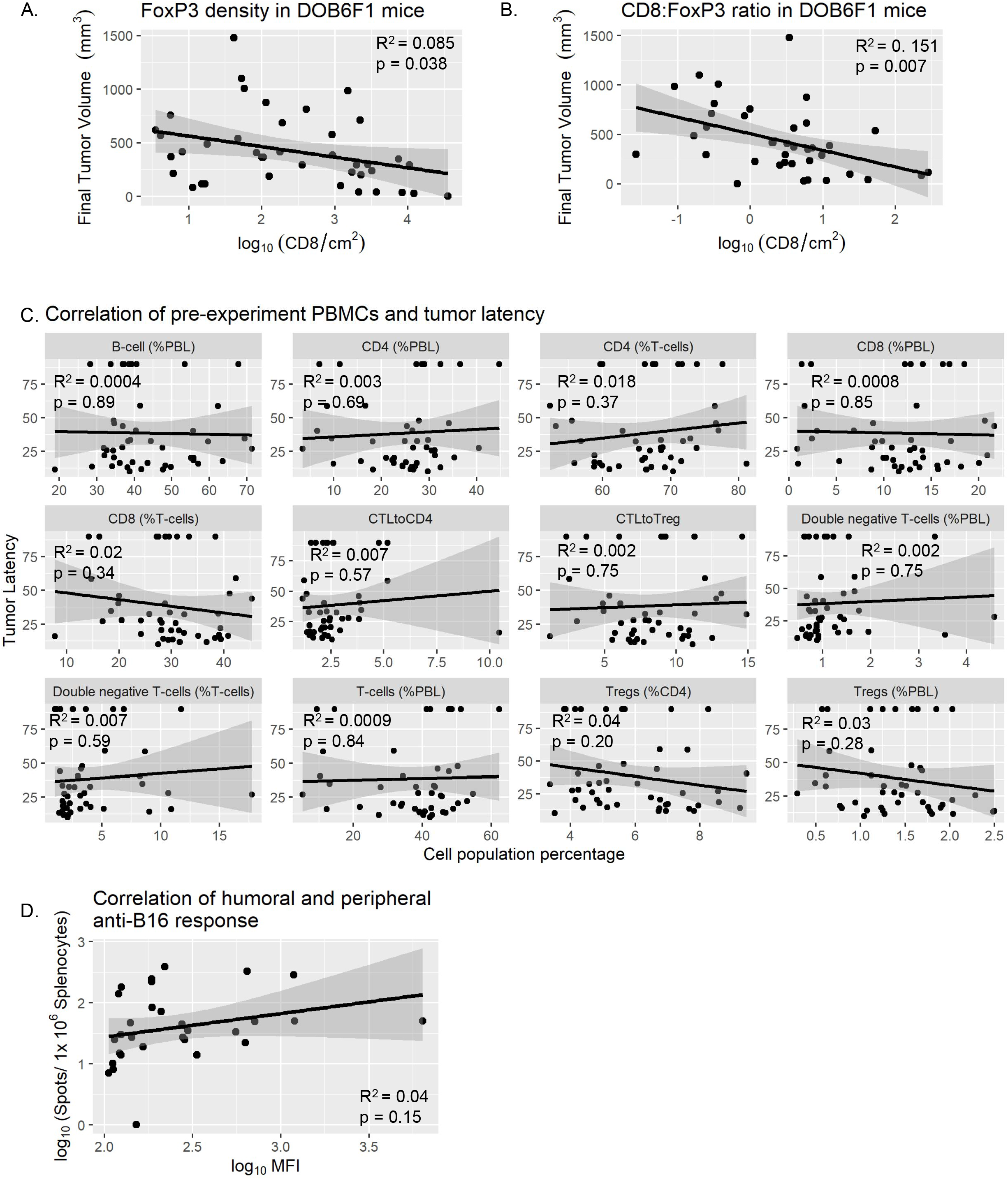
Tumor outcomes correlated against, IHC and PBL populations and correlation of humoral and cellular anti-tumor response **A)** Infiltration of FoxP3 measured by IHC plotted against final tumor volume. Correlation was evaluated using Pearson’s correlation. **B)** Ratio of CD8 and FoxP3 infiltrates measured by IHC plotted against final tumor volume. Correlation was evaluated using Pearson’s correlation. **C)** Correlation of pre-existing PBMC populations and tumor latency determined by flow cytometry. **D)** Mean Fluorescence Intensity (MFI) of anti-B16F0 IgG in serum from cohort 4 plotted against ELISPOT counts from same animals. Correlation was evaluated using Pearson’s correlation.

**Fig S2.**
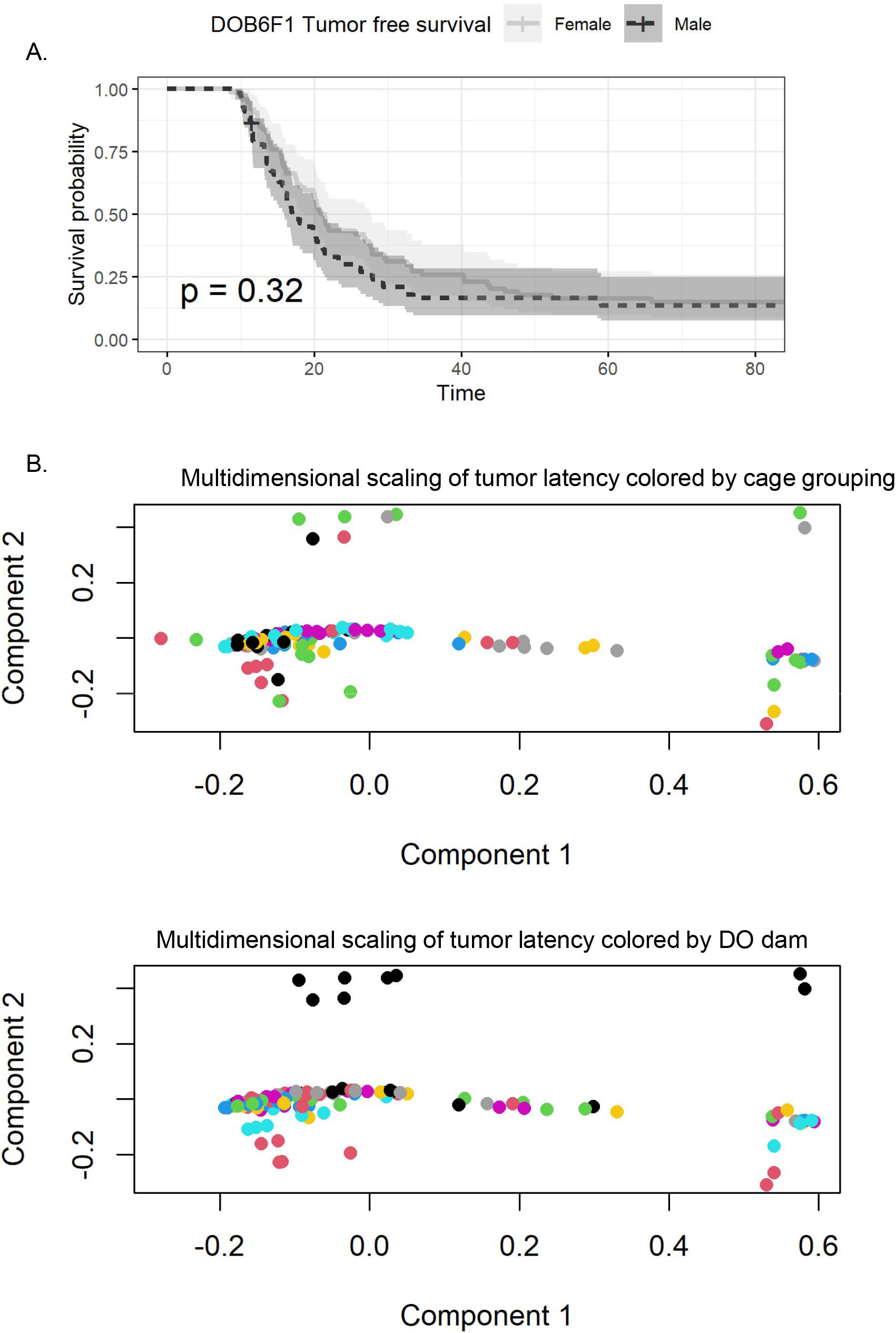
Association of sex, cage, and siblingship to tumor latency. **(A)** Tumor-free survival risk based on sex in treated DOB6F1 mice was evaluated by Kaplan-Meier estimate. **(B)** Plot of tumor latency against DO founder contribution from DO dam. Correlation evaluated using Pearson’s correlation. **(B)** Multidimensional scaling from dissimilarity matrix created using the Gower distance metric of tumor latency with cage grouping and mom as factors.

**Fig S3.**
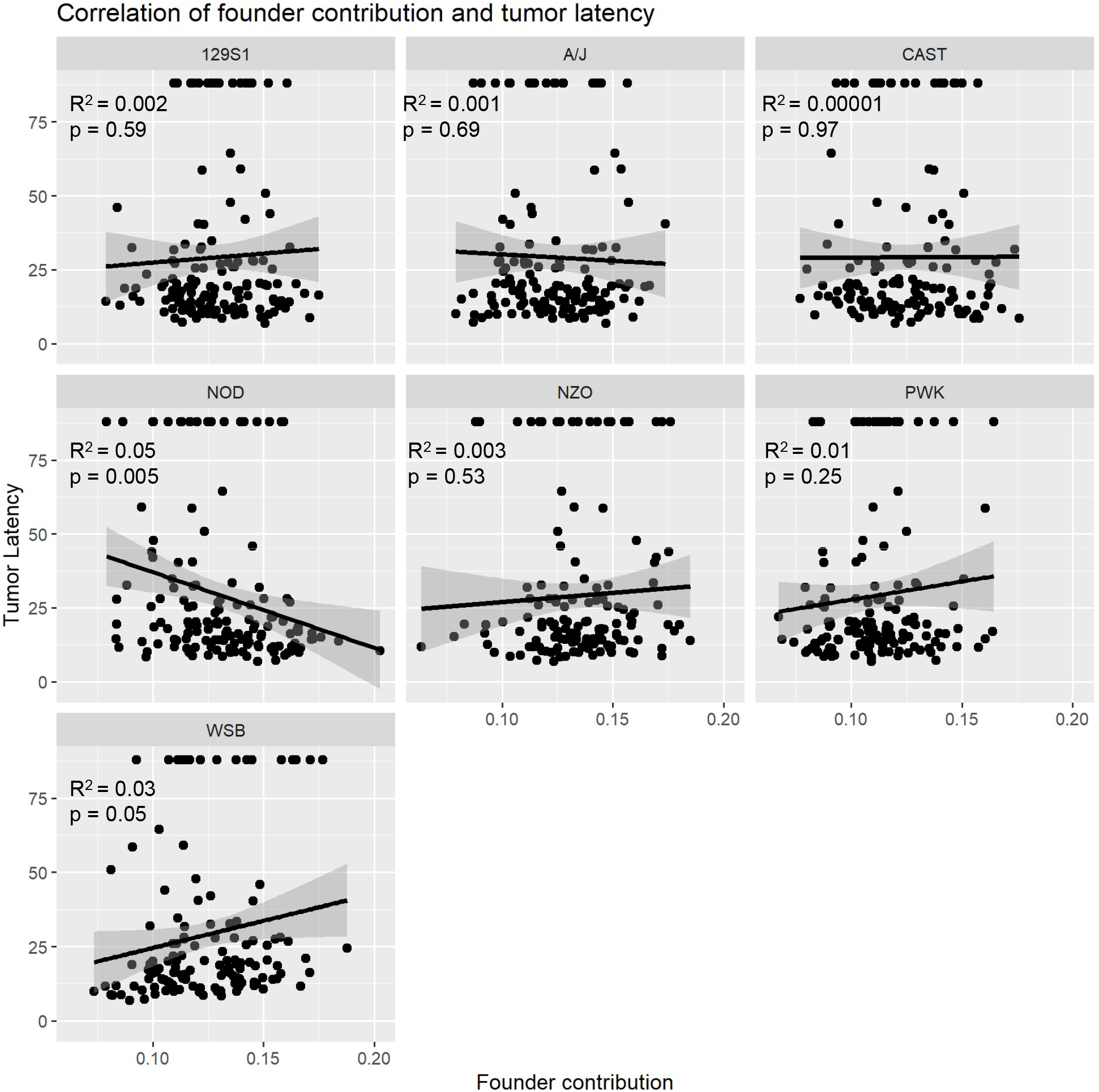
Correlation of founder proportion and tumor latency in DOB6F1 mice. **(A)** Plot of founder contribution from DO dam against tumor latency. Correlation evaluated using Pearson’s correlation.

**Fig S4.**
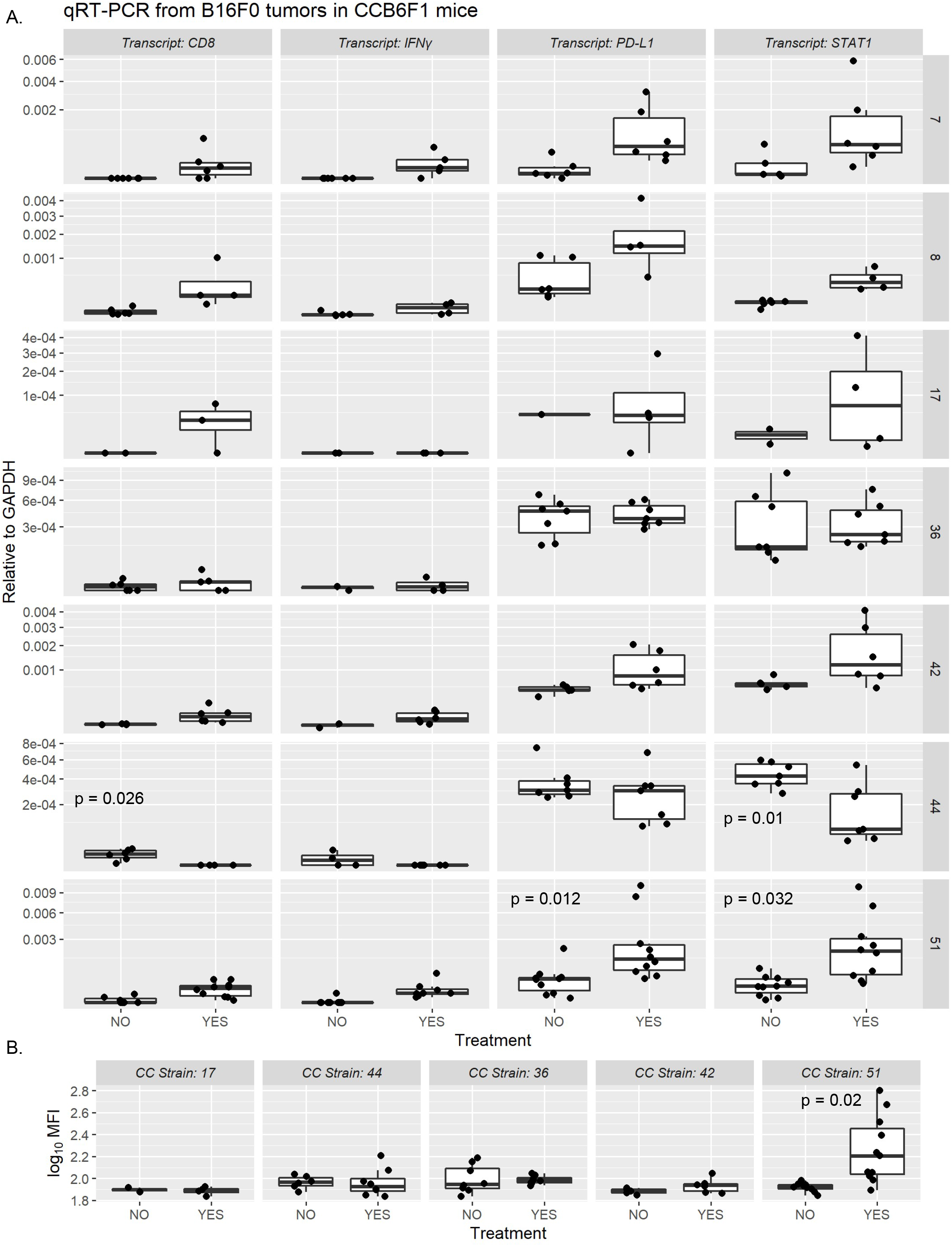
qRT-PCR from flash frozen tumors from CCB6F1 mice with and without ICI and anti-B16 antibody from select CCB6F1 mouse strains. **A)** qRT-PCR of CD8, IFNγ, and downstream IFNγ response genes PD-L1 and STAT1 relative to GAPDH from flash frozen tumors in ICI treated and untreated CCB6F1 mice. Nondetects in samples were set to a Ct of 55 to reduce bias. Significance is determined by Student’s T-test. **B)** Mean Fluorescence Intensity (MFI) of anti-B16F0 IgG in serum of CCB6F1 mice measured by flow cytometry. Statistical significance determined via Student’s T-test.

**Fig S5.**
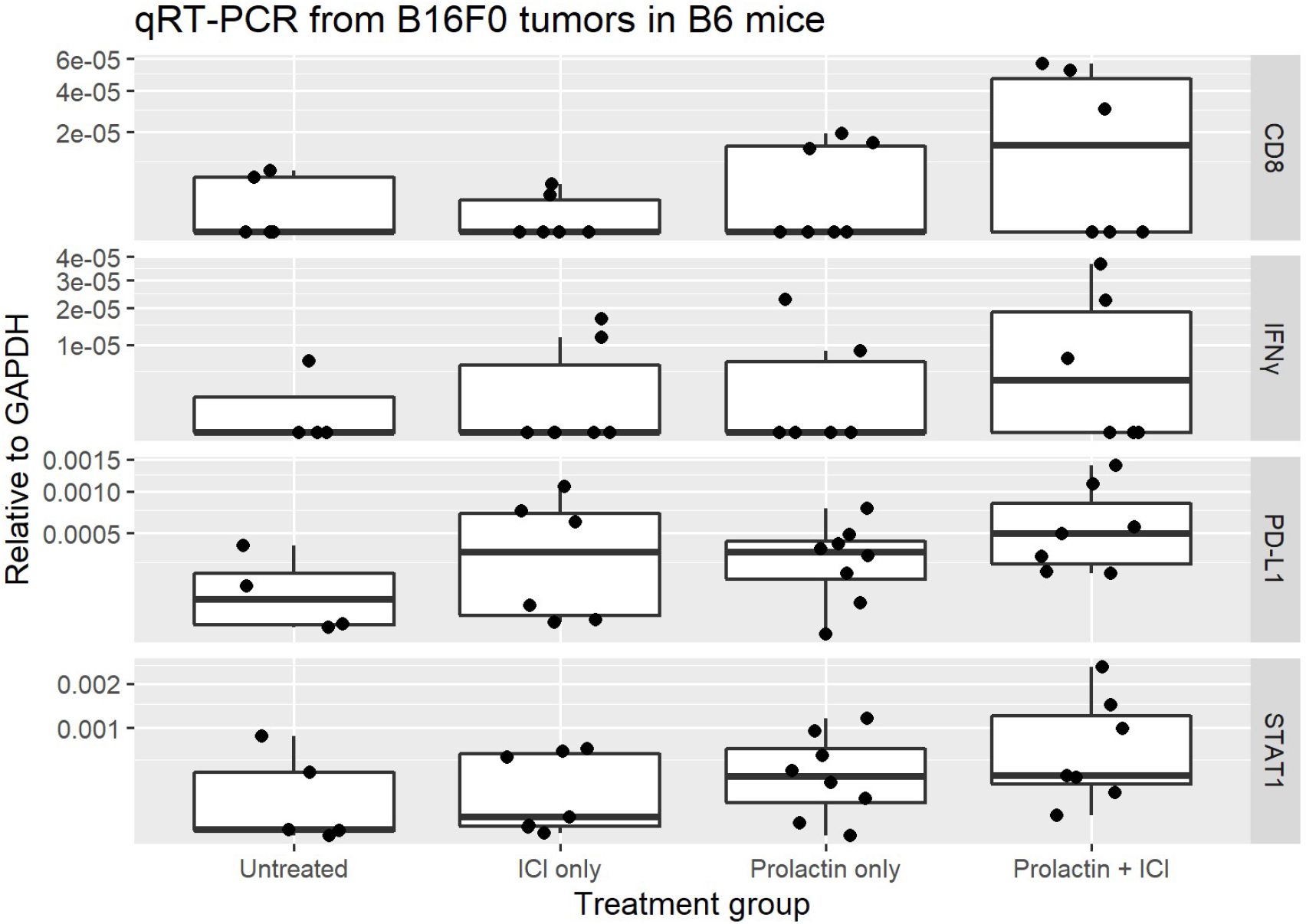
qRT-PCR from flash frozen tumors in B6 mice receiving ICI, prolactin, or combined ICI + prolactin. **A)** qRT-PCR from flash frozen tumors from **(Figure 6)** for CD8, IFNγ, and downstream IFNγ response genes PD-L1 and STAT1.

**Fig S6.**
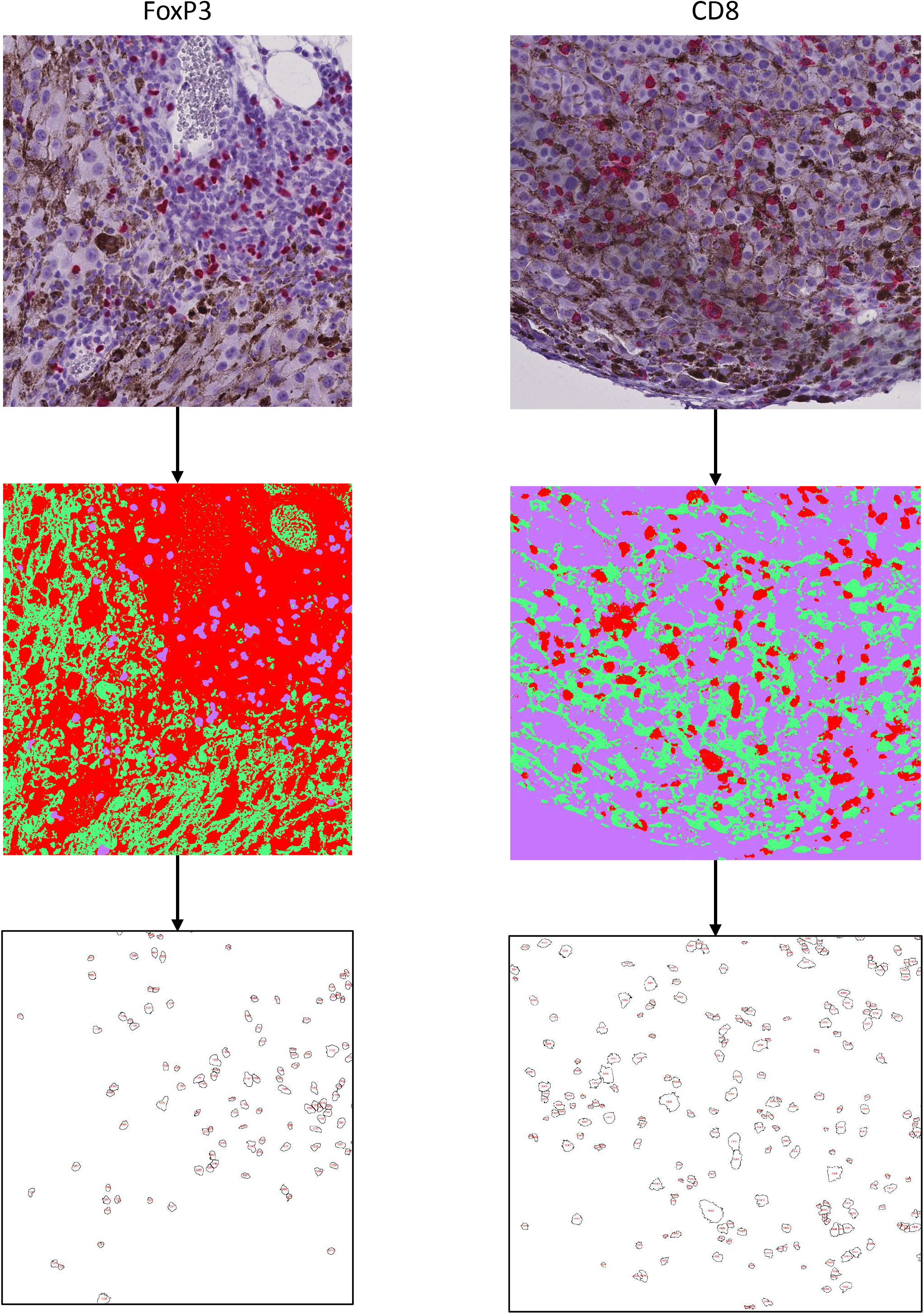
FoxP3 and CD8 IHC quantitation of representative sections. Weka segmentation for counting positive IHC in whole slide scans. Top image shows representative section of an original slide scan. Second image shows image after classifying pixels into tumor (purple), melanin (green), and positive staining (red). Final image shows resulting identified cells that were counted.

**Fig S7.**
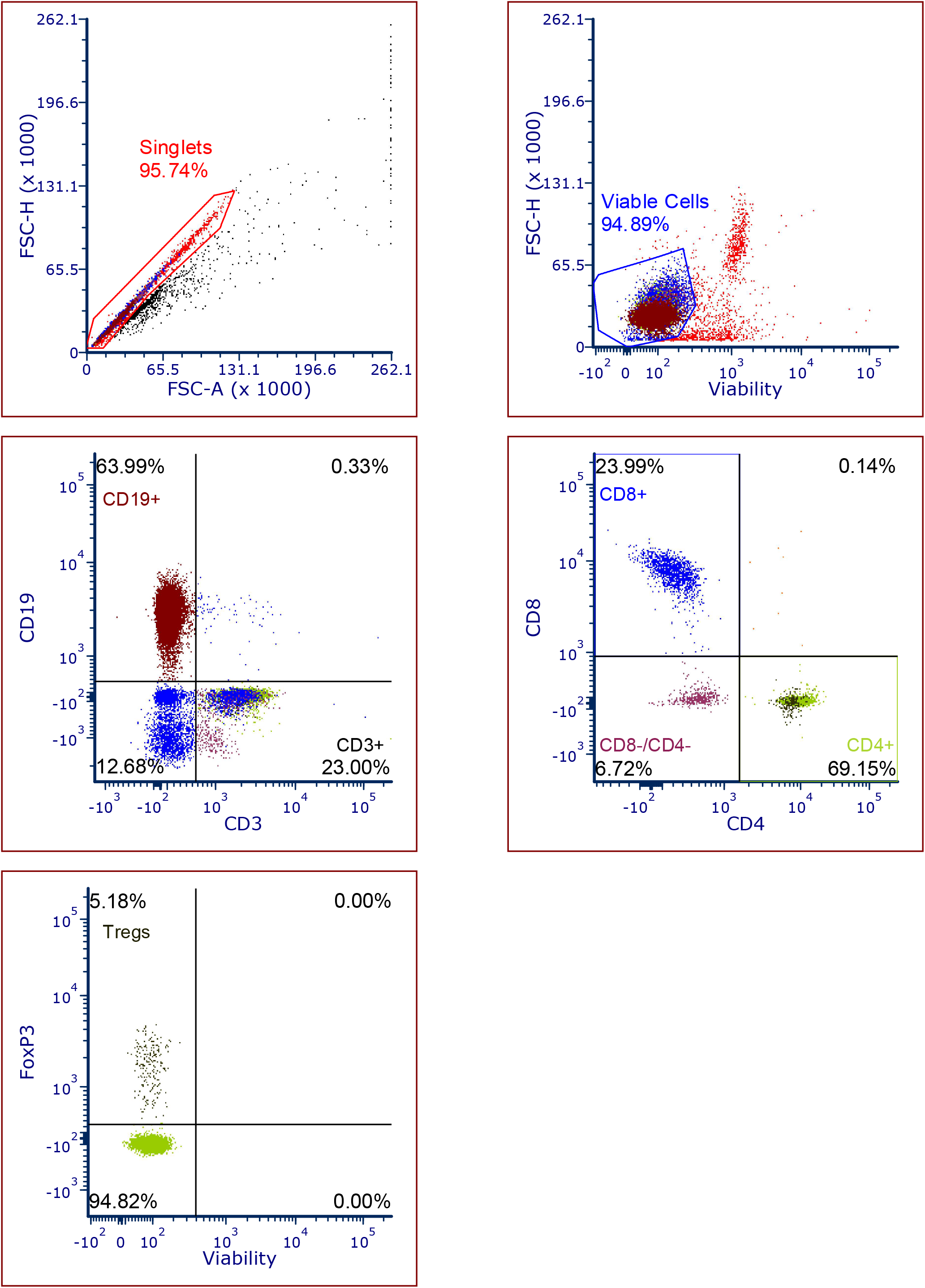
Example gating strategy of PBLs. Gating strategy showing selection of cell types used to correlate PBL populations to tumor latency.

